# Proteomics Defines Shared and Divergent Alterations in the Right Atrium and Right Ventricle in Porcine Right Heart Failure

**DOI:** 10.64898/2026.01.08.698419

**Authors:** Sally E. Prins, Jacob Sternbach, Jenna B. Mendelson, Todd Markowski, LeeAnn Higgins, John P. Carney, Walt Tollison, Gaurav Choudhary, Kurt W. Prins

## Abstract

Right heart failure is due to both right atrial and right ventricular dysfunction. While the two chambers are distinct, the proteomic response to pressure overload is undefined. Here, we used quantitative proteomics to evaluate changes in both chambers in pulmonary artery banded pigs. We found common alterations in oxidative phosphorylation, ribosome regulation, integrin-mediated adhesion, and glycolysis enzymes. However, the right atrium had more pronounced deficits in the TCA cycle and branched chain amino acid metabolism. Thus, these data highlight metabolic pathways that could be evaluated to selectively enhance right atrial function.

---

Right heart failure (RHF) is a leading cause of morbidity and mortality in pulmonary arterial hypertension (PAH), and we currently lack effective therapies to combat this deadly condition^1^. Proper right heart function is dependent on the synergistic actions of the right atrium (RA) and the right ventricle (RV) for the circulation of deoxygenated venous blood through the pulmonary vasculature and to ultimately fill the left ventricle. Recent studies demonstrate right atrial dysfunction is an emerging risk factor for heightened mortality in PAH^2^, but the underlying pathobiology of right atrial dysfunction is poorly understood. In addition, a comparison of the chamber specific alterations in the diseased right atrium and right ventricle is lacking, which limits our ability to effectively treat both chambers to combat RHF.

Control and pulmonary artery banded (PAB) pigs and their cardiac MRI examinations were previously described^3,4^. Crude mitochondrial enrichments (Abcam) from RA (*n*=4 control and *n*=4 PAB) and RV (*n*=4 control and *n*=4 PAB) tissues were generated and subjected to TMT16plex proteomics at the University of Minnesota Center for Metabolomics and Proteomics. Protein abundances were determined using Proteome Discoverer v3.0. Calculated protein abundances were correlated with RV ejection fraction or RA emptying fraction to determine which proteins were associated with right heart chamber function. Proteins with correlational values of |*r*|>0.5 with either RA emptying fraction or RV ejection fraction were subjected to Wiki pathway analysis using Shinygo v0.85. Relative proteomic pathway scores were calculated by summing the abundances of all proteins detected in each pathway and values were then normalized to the average of the four-control RA or the four RV samples, respectively. Statistical analysis was performed using GraphPad Prism v 10.5.0. Unpaired *t*-test or Mann-Whitney test compared the differences between two groups pending normality as determined by Shapiro-Wilk test. Raw proteomic data are available at zenodo.org, doi:10.5281/zenodo.17085369.

Hierarchical cluster analysis revealed PAB induced proteomic alterations in both the RA and RV (**Figure A**). Next, we performed correlational analysis of protein abundances with cardiac-MRI determined measures of RA and RV function. In the RA, Wiki pathways associated with enhanced RA emptying fraction included the tricarboxylic acid (TCA) cycle, branched chain amino acid (valine, leucine, and isoleucine, BCAA) metabolism, and oxidative phosphorylation (**Figure A**). In the RV, Wiki pathways associated with higher RV ejection fraction were dominated by oxidative phosphorylation, which suggested slightly different mechanisms of metabolic remodeling occurred when comparing the RA to the RV (**Figure A**). On the other hand, pathways associated with impaired chamber function were more concordant as ribosomal, integrin-associated cellular adhesion, and glycolytic pathways were identified in both RA and RV tissue (**Figure A**).

Then, we compared whole pathway regulation in both chambers. Although not always statistically significant, both the diseased RA and RV exhibited downregulation of oxidative phosphorylation enzymes and upregulation of ribosomal, glycolysis, and integrin associated cellular adhesion pathways (**Figure B**). Interestingly, only the RA exhibited downregulation of the TCA cycle and BCAA catabolism proteins (**Figure B**).

**Figure.**
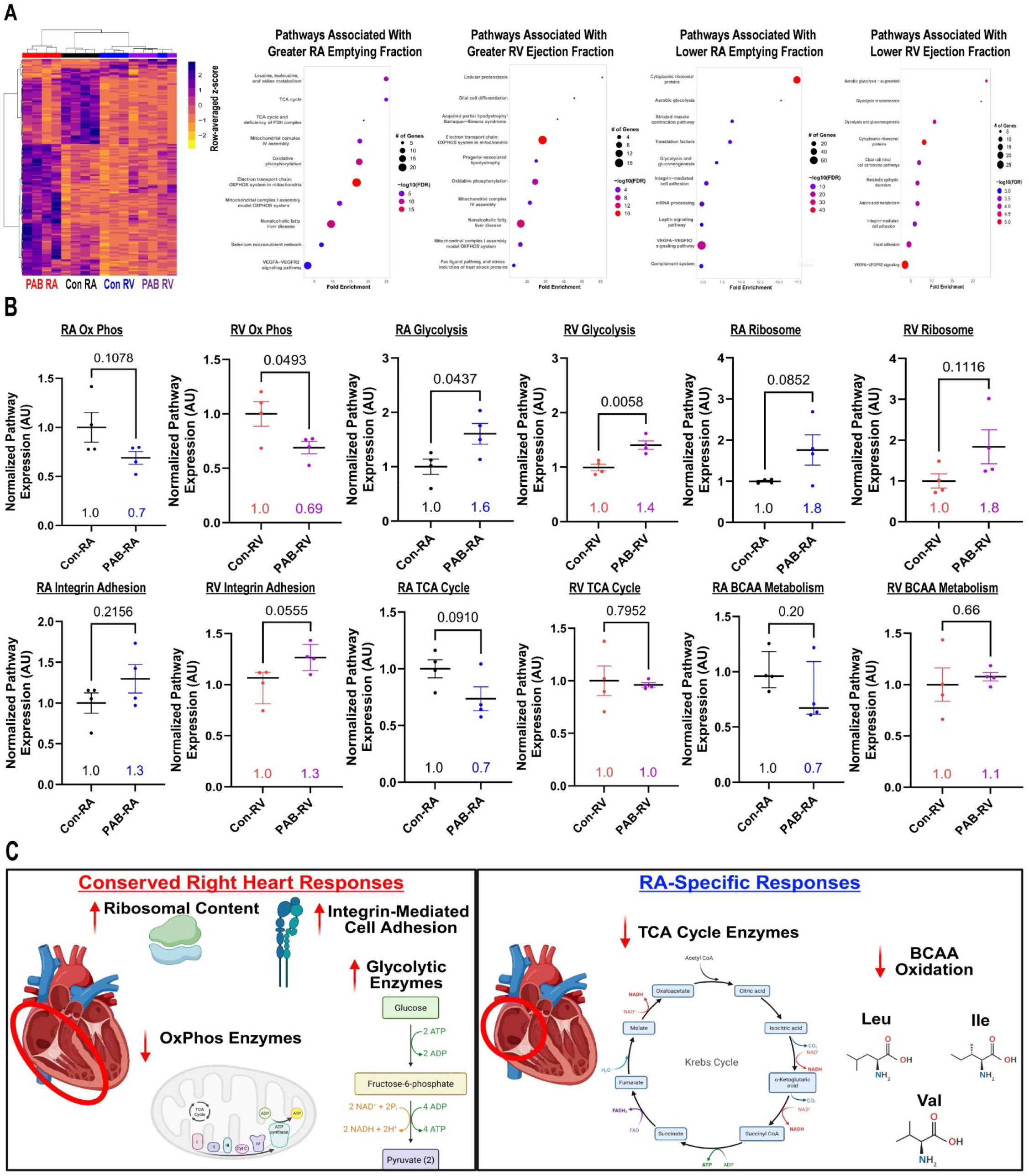
Proteomics analysis defined shared and divergent pathway associations with chamber-specific function in porcine right heart failure. (A) Hierarchical cluster analysis of proteomics evaluation demonstrated alterations in both the RA and RV following PAB. Wiki pathway analysis of proteins that were enriched with greater RA emptying fraction or RV ejection fraction (left). Wiki Pathway analysis of proteins that were enriched with lower right atrial emptying fraction and right ventricular ejection fraction (right). (B) Relative pathway score of oxidative phosphorylation, glycolysis, ribosomes, integrin-mediated cell adhesion, TCA cycle, and BCAA metabolism. *p*-values determined by unpaired *t*-test or Mann Whitney test. Numerical values represent mean or median value. Data on graphs are presented as mean plus standard error of mean or median plus interquartile range (C) Summary of conserved alterations in both the RA and RV(left) and RA-specific disruptions in pulmonary artery banded pigs (right).

In conclusion, our data nominates pathways that may contribute to right heart failure via both RA and RV compromise with shared alterations in oxidative phosphorylation, glycolysis, ribosomes, and integrin-mediated adhesion. In addition, we identify the TCA cycle and BCAA metabolism as RA-enriched alterations that could serve as targets to more specifically improve RA function. In agreement with this hypothesis, a mutagenesis screen identified branched chain amino acid transaminase 2, an enzyme key for BCAA catabolism, as an important gene for regulating cardiac arrhythmias *in vivo*^*5*^. Moreover, excess BCAAs induce calcium handling abnormalities and pro-arrhythmic events *in vitro*^*5*^. Perhaps augmenting BCAA catabolism would enhance RA function via both rhythm and contractile mechanisms.

Our findings have important limitations including use of only young, castrated male pigs, and our findings are only hypothesis generating. Whole tissue extracts were used in our proteomics analysis, so we lack resolution to determine which cell types underlie the overall changes in the proteins reported. The analysis of total pathway engagement did not result in significantly different changes in many instances, but not all proteins in each pathway were dysregulated as only components of each pathway were associated with chamber function. Finally, future studies are required to determine if pathway activation or suppression is maladaptive or compensatory in each chamber. It is plausible that greater abundances of ribosomes and integrin-adhesions are protective to promote protein synthesis required for hypertrophy and reinforce cell-cell interactions due to heightened force generating requirements. Thus, negating these changes could have deleterious effects.

